# Variable inhibition of different Legionella species by antagonistic bacteria

**DOI:** 10.1101/2024.11.27.625680

**Authors:** Alessio Cavallaro, Silke Probst, Tobias Duft, Max Rieder, Oliver Abo El Fateh, Josch Stricker, Marco Gabrielli, Serina Robinson, Frederik Hammes

## Abstract

The genus Legionella includes opportunistic pathogens inhabiting engineered aquatic ecosystems, where managing their presence and abundance is crucial for public health. In these environments, Legionella interact positively or negatively with multiple members of the microbial communities. Here, we identified bacteria and compounds with Legionella-antagonistic properties. We isolated 212 bacterial colonies from various water sources in Switzerland and screened them for their ability to inhibit one reference strain of L. pneumophila. Ten selected antagonistic isolates were subsequently tested with spot-on-lawn-assays for inhibition towards seven environmental and two clinical isolates of Legionella, representing different species and strains. The antagonists produced highly variable inhibition patterns, highlighting distinct differences in susceptibility among Legionella species, and even strains. Only three isolates, all identified as Pseudomonas lurida, inhibited all Legionella species. Furthermore, we analysed the genomes of the antagonistic bacteria, and identified genes for several probable inhibitory compounds. We specifically found the gene cluster for the biosurfactant viscosin to be uniquely encoded by three Pseudomonas lurida isolates. This compound was subsequently detected in the supernatant of co-cultures inoculated with the antagonists and Legionella. This study provides new insights on the ability of aquatic microorganisms to compete with Legionella in controlled laboratory settings. It also highlights the diversity across and within Legionella species in their resistance to external antagonistic stress, and confirms the anti-Legionella activity of selected biosurfactants. These results can contribute to the understanding of how different species inhabit separate niches in the environment, and expand the discussion around alternative Legionella mitigation strategies.

## 1. Introduction

Bacteria belonging to the genus Legionella are waterborne opportunistic pathogens and aetiological agents of Legionnaires’ disease, one of the most reported water-associated illnesses and with increasing incidence worldwide (1–3). Legionella pneumophila accounts for more than 90% of these cases (1, 4), yet the broad Legionella genus comprises to date more than 70 species officially recognized (5), of which approximatively half have been already described as human pathogens (6, 7). The ecology of these non-pneumophila species is often less studied, although relevant given the fact that many of them are co-occurring in the environment (8). Legionella are typically found in engineered aquatic ecosystems such as building plumbing or cooling towers (2, 9), where their presence is mostly managed through chemical and physical disinfection. However, Legionella often persist in the environment due to their physiology and ecological characteristics, such as the protection offered by biofilms and protists (3, 10). In fact, Legionella live embedded in complex biofilms, where they interact with multiple prokaryotic and eukaryotic organisms creating specific environmental niches (10).

In recent years, several studies have investigated the action of various compounds of biological origin to inhibit the growth of Legionella. Many of these discoveries have been well summarized by Berjeaud and coworkers (11), who divided the identified compounds in five classes, including proteins (e.g., apolipophorin III and lactoferrin isolated from eukaryotic organisms), protein-derived peptides of synthetic origin, antimicrobial peptides (e.g., warnericin RK isolated from Staphylococcus warneri), essential oils, and biosurfactants (e.g., surfactin isolated from Bacillus subtilis). However, these were isolated in diverse biological contexts not specifically linked to Legionella’s natural habitats, hence some of these mechanisms may not necessarily occur in natural environments.

Targeted competition experiments have explored the anti-Legionella potential of microorganisms isolated directly from Legionella’s natural aquatic habitats. For example, Guerrieri and coworkers (12) tested 80 bacteria isolated from tap water and found that 37 out of 80 were active against at least one of 26 strains of L. pneumophila. Similarly, Corre and coworkers (13) reported the antagonistic activity of 178 bacteria collected in their study towards one strain of L. pneumophila, while Paranjape and coworkers (14) described seven isolates from cooling towers being able to inhibit multiple strains of L. pneumophila, and detected the presence of gene clusters potentially encoding for molecules potentially responsible for the inhibition. In most of these cases, bacteria belonging to the genus Pseudomonas, known to produce diverse secondary metabolites (15), were detected as the main antagonists.

While these competition studies provide valuable information on the identity and biological activity of several aquatic bacteria towards Legionella, the actual inhibitory mechanisms are rarely identified, with the exception of biosurfactants produced by Pseudomonas strains reported by Loiseau and coworkers (16). Moreover, the majority of antagonism studies were conducted using only L. pneumophila as the target strain, and therefore little is known about how other Legionella species respond to antagonism.

In the present study, we expand on previous work by testing the inhibitory activity of multiple bacterial isolates obtained from different water sources in Switzerland through spot-on-lawn assays. Importantly, we specifically focused how different Legionella species respond to the antagonistic activity. We moreover investigated the antagonistic isolates for the presence of potential biosynthetic gene clusters encoding specific secondary metabolites, as the probable cause for the inhibition, and then detected their presence through liquid chromatography and mass spectrometry. Finally, we discuss our results in a broader environmental perspective, highlighting the role of different Legionella species in establishing distinct ecological niches.

## 2. Materials and Methods

### 2.1. Antagonist isolation

Water samples were collected from various locations and sources in Switzerland (Supplementary Table S1) in 100 mL Schott bottles and 1 mL of the same sample was subsequently plated onto Lysogeny Broth (LB) and R2A agar plates (200 µL per plate) that were incubated at 30 °C for five days. Twelve colonies per sample were picked based on distinct morphological properties and re-plated for isolation. The isolates were subsequently cultured in liquid medium overnight and glycerol stocks (25% V/V) were prepared for storage at -80 °C.

### 2.2. Legionella **strains**

A reference strain of Legionella pneumophila (DSM7513) was obtained from the German collection of microorganisms (DSMZ). The other Legionella isolates used in this study were obtained from the Swiss National Legionella Reference Centre (CNRL) located in Bellinzona (CH). The strains were selected among confirmed human pathogens and originated from either environmental water samples or clinical samples. A full list of the provided strains can be found in Table 1.

**Table 1.**
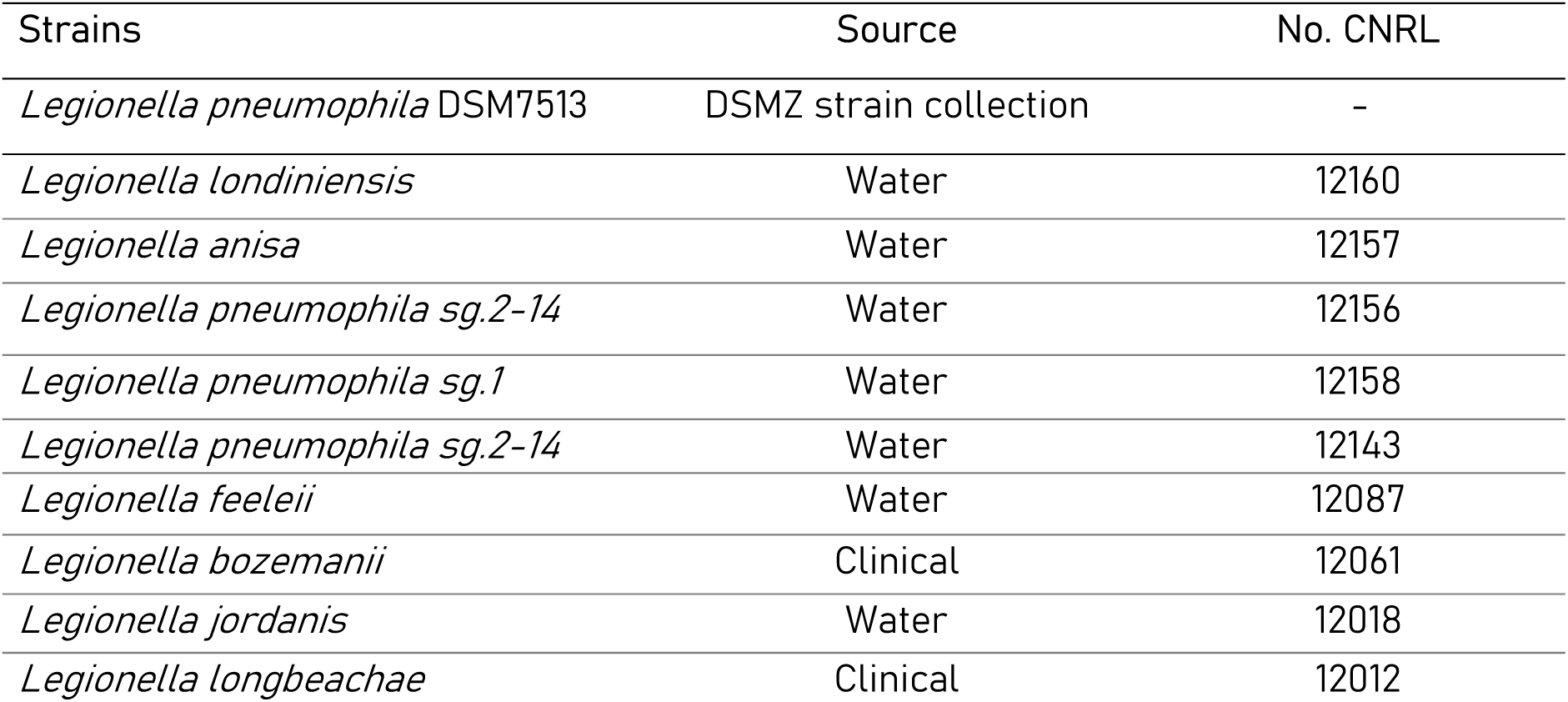
List of Legionella strains used in this study.

### 2.3. Spot-on-lawn assay

Spot-on-lawn experiments were performed to evaluate the inhibition of Legionella. Briefly, Legionella strains were grown for 72 h on Buffered Charcoal Yeast Extract (BCYE) plates, and a colony was then transferred into Buffered Yeast Extract Broth (BYEB). After 72 h, the total cell count of Legionella in the culture was quantified using a CytoFLEX (Beckman Coulter, Brea, USA) flow cytometer in 250 µL aliquots stained using SYBR® Green I (SG, Invitrogen AG, Basel, Switzerland; 10,000x diluted in Tris buffer, pH 8). Stained cells were incubated for 15 minutes at 37 °C prior to analysis (Prest et al., 2013). After diluting the cultures to a concentration of 10^4^ cells/µL, 100 µL was inoculated on a BCYE agar plate and distributed with a cotton swab in order to form a dense lawn. To test the inhibitory activity of the bacterial isolates, fresh cultures were grown overnight and diluted to the same concentration as the Legionella strains. 10 µL of the diluted cultures were spotted onto the agar plates previously inoculated with the Legionella lawn and the plates were incubated at 30 °C for 72 h. To test the inhibitory properties of the HPLC fractions (details in Section 2.5), these were diluted in methanol after being dried. Oxoid™ Blank Antimicrobial Susceptibility discs were soaked with 50 µL of each fraction and placed onto BCYE agar plates previously inoculated with the Legionella lawn (as above). An inhibitory activity was detected when a zone of non-visible growth was observed in the Legionella lawn around the colony or the susceptibility disc. All the experiments were conducted in triplicate, and the plates were imaged using a camera set-up that allows for reproducible imaging. The inhibition was measured as the diameter of the inhibition zone, normalized by accounting for the colony size of the antagonist. Examples of the spot-on-lawn assays conducted with bacterial isolates or discs are provided in Figure 1 and Supplementary Fig. S1.

**Figure 1.**
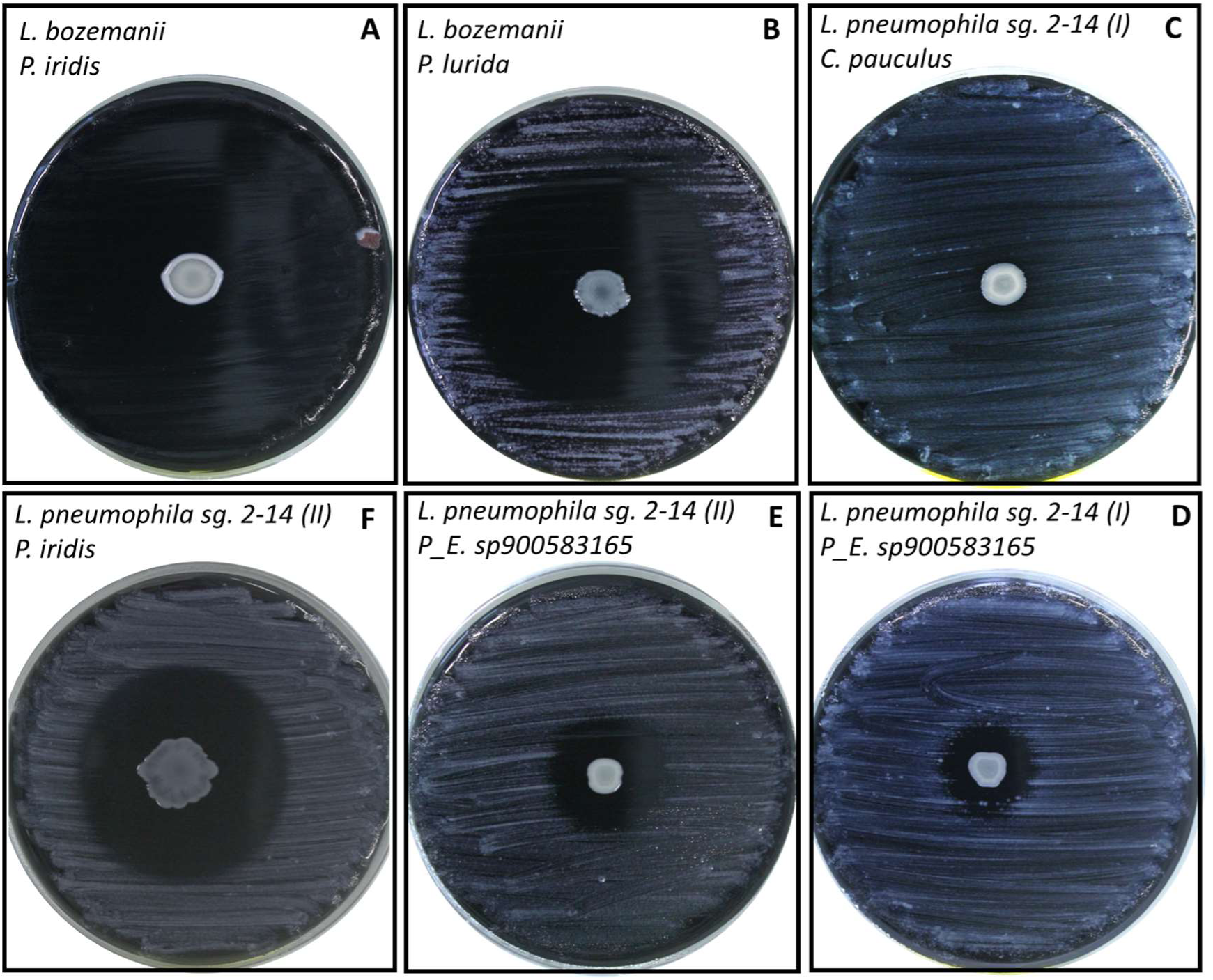
Examples of the spot-on-lawn experiments with multiple Legionella species and selected antagonistic isolates, showing variable visible outcomes. The top left corner provides the description of the Legionella strain used (top) and the antagonist (bottom). (A) Total inhibition of one Legionella species that is (B) only partially inhibited by a different antagonist. (C) No visible inhibition for a different Legionella strain, while (D) the same strain is mildly inhibited by a different antagonist. Different degrees of inhibition for the same Legionella species, when subjected to the action of different antagonists in E and F.

### 2.4. Whole genome sequencing and bioinformatics

The genomic DNA of ten selected antagonistic bacteria was isolated using the Qiagen DNeasy Blood and Tissue extraction kit. Whole Genome Sequencing was performed in service by SeqCenter in Pittsburgh, PA (USA). Sample libraries were prepared using the Illumina DNA Prep kit and IDT 10bp UDI indices, and sequenced on an Illumina NextSeq 2000, producing 2x151bp reads. Demultiplexing, quality control and adapter trimming was performed with bcl-convert (v3.9.3). Genomes were then assembled using the software SPAdes (v3.15.5) (17), and further quality-controlled for contamination and chimerism using the software CheckM2 (v1.0.2) (18) and GUNC (v1.0.6) (19). Taxonomy was assigned using GTDB-Tk (v2.3.2) through the function “classify_wf” (20). Genome mining for secondary metabolites was performed using the software antiSMASH (v7.0) (21) using the default parameters. All plots and tables were generated in R (v4.3.1) and R studio (v2023.06.0+421) using the packages ggplot2 (v3.4.2) and kable (v1.4.0).

### 2.5. Liquid-liquid extraction and HPLC fractionation

Co-cultures of L. jordanis and P. lurida (I) were prepared inoculating pre-grown bacteria diluted to a concentration of 10^4^ cells/µL, in a ratio Legionella:Pseudomonas of 7:3. The co-culture was incubated for 72 h at 30°C in an orbital shaking incubator. After incubation, the culture was centrifuged using a Multifuge X Pro Series (ThermoScientific) at 10’000 g for 15 minutes and the supernatant separated from the pellet. The supernatant underwent liquid-liquid extraction with ethyl acetate. Briefly, 1.5 times volume of ethyl acetate was added to the supernatant in a separation funnel, and the mixture shaken thoroughly in order for the compounds to migrate to the organic phase. The extract was then transferred for subsequent analysis. The procedure was repeated three times using the same supernatant. The extracted organic phase was evaporated to dryness using a rotary evaporator and stored at -20°C until further use. The frozen extract obtained from the liquid-liquid extraction was reconstituted using 2 mL of methanol, and the precipitate was dissolved in a sonication bath. The sample was then centrifuged at 14’000 rpm for one minute in order to separate any particles from the liquid to inject in the HPLC machine. The setup for the reverse phase HPLC consisted of a HPLC machine (Dionex UltiMate 300 Pump, ThermoScientific) and a column (C18(2), 250 x 21, Luna 5). The column was connected to a detector (Dionex UVD 340U Detector, Gynkotek) and a fraction collector (Model 2128, BIO-RAD). The collector switched between glass vials every minute, collecting approximately 10 mL of solution in each glass vial. The program for the liquid phase during the run was as follows: (I) Increase of acetonitrile from 5% to 100% within 15 minutes; (II) 15 minutes of 100% acetonitrile; (III) Decrease of acetonitrile from 100% to 5% within 5 minutes. Before starting the program, 1 mL of the 2 mL sample was added into the HPLC system using a syringe. A total of 31 fractions were collected. Two additional fractions were collected during the wash cycle, since an additional two peaks were detected. The fractions were then evaporated using a vacuum pump (Vacuum Pump V-600, Büchi). The concentrated fractions were dissolved in 1 mL of methanol and stored until further use.

### 2.6. Liquid Chromatography – Mass Spectrometry (LC-MS)

The supernatant of the L. jordanis – P. lurida co-culture (above), along with supernatant from mono cultures of L. pneumophila sg. 1, L. jordanis and P. lurida (I, II) and the fractions that showed inhibition of Legionella in spot-on-lawn assays, were subjected to LC-MS to verify the presence of viscosin. All the supernatants underwent liquid-liquid extraction as described in Section 2.5. The samples were run through a HPLC system (Ultimate 3000, Dionex) connected to a column (2.6 μm XB-C18 150 x 4.6 mm, Phenomenex Kinetex). A mass spectrometer system (LTQ Orbitrap XL, Thermo ScientificTM) was coupled to the HPLC, resulting in HPLC-HRESI MS analytical analysis MS spectra. A specific program was used for each sample type using two solvents. Solvent A consisted of H2O and 0.1% formic acid, while solvent B consisted of acetonitrile and 0.1% formic acid. The injection volume of 10 μL was used for each sample with a flowrate of 0.7 mL/min. The solvent programs are shown in Table 2.

**Table 2.**
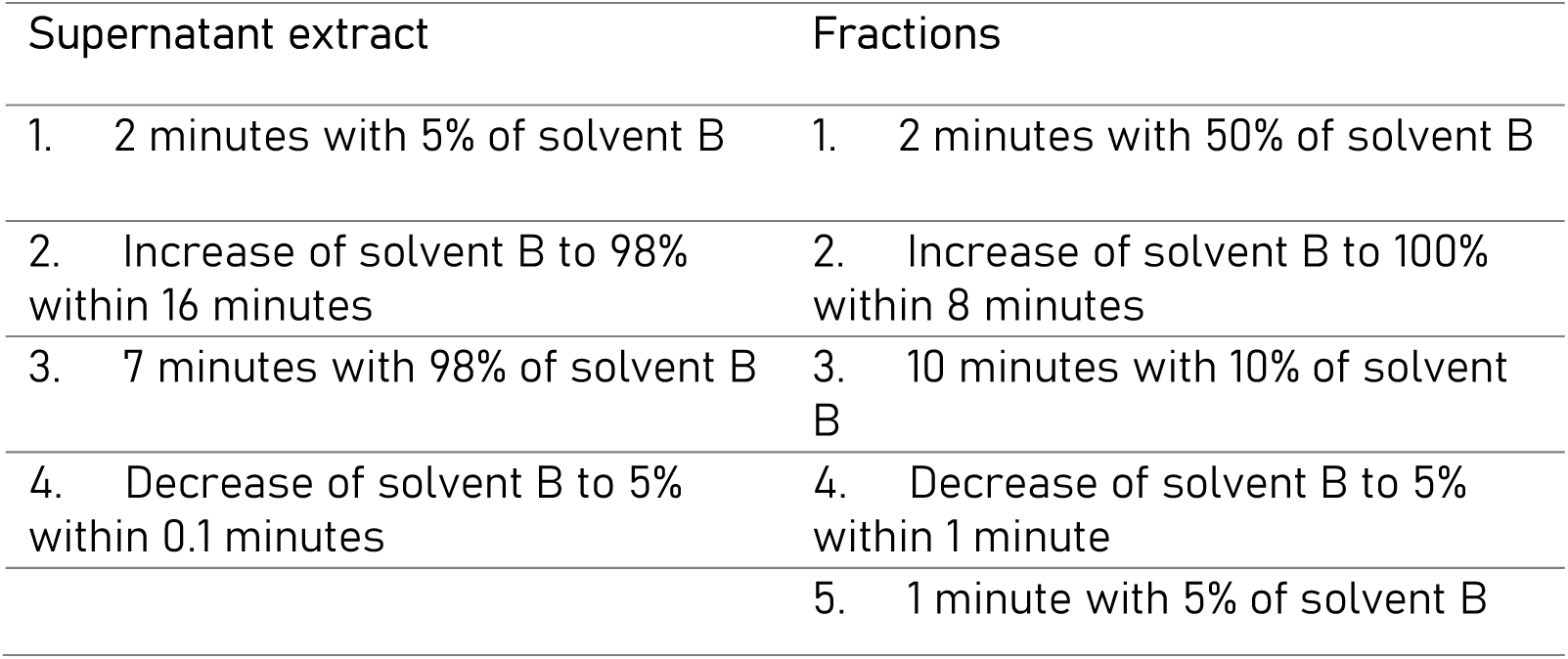
Solvent programs used in LC-MS.

## 3. Results

### 3.1. Isolation of the antagonists and inhibition of multiple Legionella species

A total of 212 isolates from 10 sources were initially screened for their ability to inhibit a reference strain of L. pneumophila DSM7513 using spot-on-lawn assays. In total, 34 isolates displayed some degree of inhibition through the formation of a clear zone in the Legionella lawn around the antagonist colony (for example, see Fig. 1). From these, 10 isolates were selected, representing the broadest morphological diversity possible among the inhibitory bacteria. Analysis of the isolates’ genomes identified six isolates from the genus Pseudomonas (three isolates belonging to the species P. lurida, one P. iridis, one P. lundensis and one Pseudomonas_E sp900583165). The rest of the isolates were classified as Brevundimonas auriantiaca (two strains), Cupriavidus pauculus (one isolate) and Sphingomonas spp. (one isolate, no information available at the species level).

However, for two of these antagonistic isolates we were unable to obtain pure cultures. Despite repeated attempts to isolate the species visibly growing together, the genomes assigned to P. lurida (III) and P. lundensis presented a high level of contamination (81.83% and 70.07%, respectively, analysis performed with CheckM2). An additional analysis performed by GUNC revealed that for P. lurida (III) 5938 genes were assigned to the genus Pseudomonas, while 4524 genes were attributed to Chryseobacterium; for P. lundensis, 4205 genes were assigned to the genus Pseudomonas, and 4814 to the genus Bacillus. Hence, the results from these two isolates should be considered in that context. The genomes of two P. lurida (I and II) isolates presented high similarity (ANI 100%; aligned fraction 99.94%). The third genome, because of the above-mentioned contamination, presented high similarity only in a portion of the genome (ANI 99.99%; aligned fraction 54.38). The two strains of B. auriantiaca presented an ANI of 99.98%, with an aligned fraction of the 90.71%.

The selected isolates were then tested against multiple environmental and clinical isolates of Legionella (Fig. 1, Table 1). The three antagonists classified as P. lurida inhibited all the Legionella species tested, but the extent of the inhibition, measured as the size of the inhibition zone, differed, indicating that the antagonists exhibit possible strain-level variability in the production of antagonistic compounds (Fig. 2). P. iridis was unable to inhibit two Legionella species (L. jordanis and L. longbeachae). Moreover, in four cases, the inhibition caused by this antagonist was total, with no detectable Legionella growth on the entire plate, strongly suggesting that volatile compounds were involved in the inhibition. Even though two antagonists were both classified as B. auriantiaca, their inhibition profiles differed: in one case, six Legionella strains were inhibited with the diameter of the inhibition zone ranging 1.53 cm to 4.64 cm; on the contrary, the second strain of B. auriantiaca only inhibited two Legionella species (L. longbeachae and L. anisa). Finally, Pseudomonas_E sp900583165 and Sphingomonas spp. only inhibited two (L. anisa and L. jordanis) and one (L. pneumophila sg.1) Legionella strains each.

**Figure 2.**
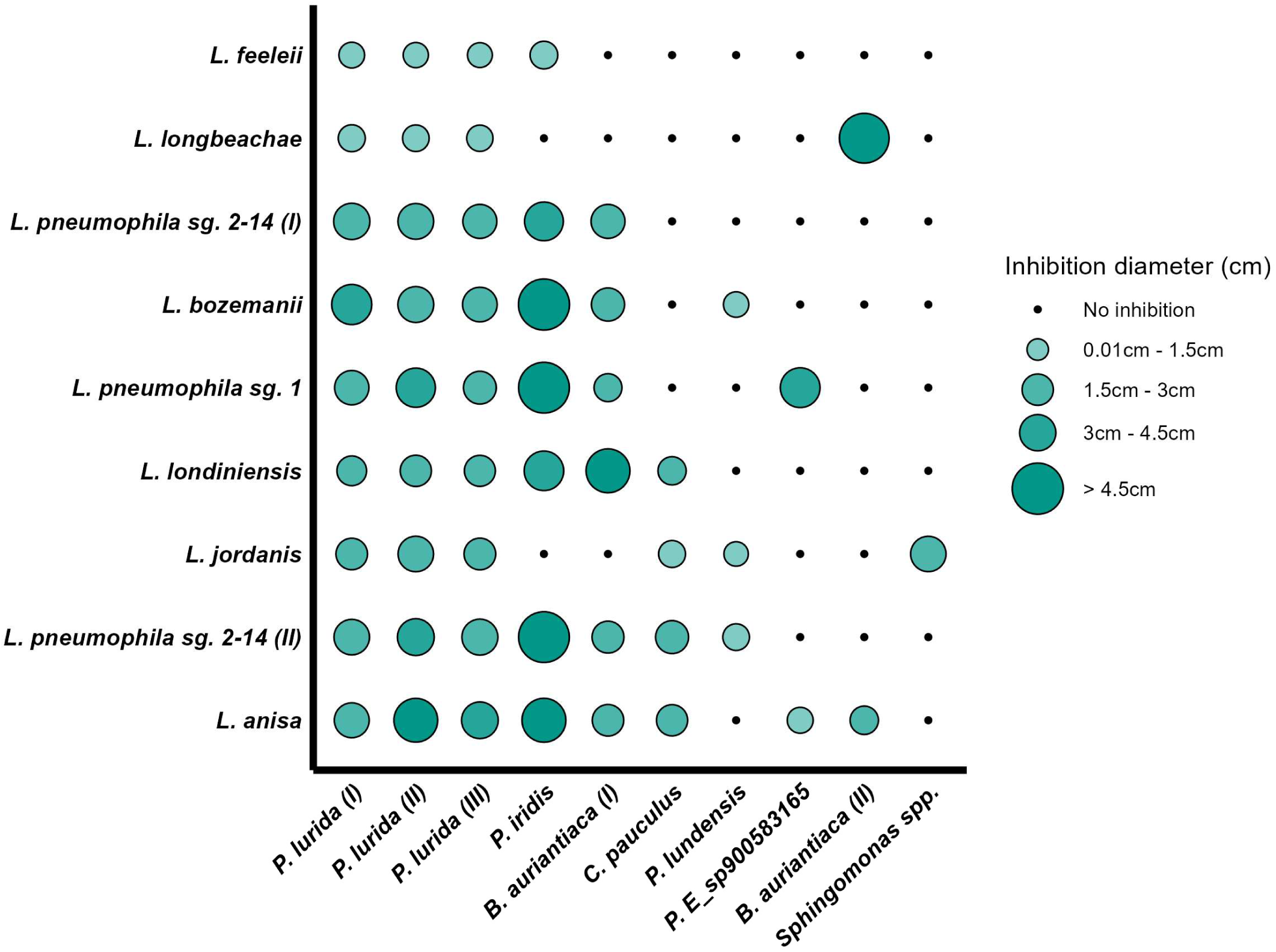
Results of the spot-on-lawn experiments in which the antagonistic isolates (displayed on the x-axis) were screened against nine clinical and environmental species of Legionella (displayed on the y-axis). A green circle indicates that an inhibition was observed on the plate, while a black point is displayed in the plot when no inhibition was detected. All the experiments were performed in triplicate; the size of the green circles is proportional to the degree of the inhibition measured as the average inhibition zone diameter, normalized for the colony size.

With respect to the Legionella strains, L. anisa was the most susceptible strain, inhibited by eight different antagonists. Interestingly, the strains of L. pneumophila displayed a different inhibition pattern: while L. pneumophila sg. 2-14 (II) was susceptible to seven antagonists, L. pneumophila sg. 2-14 (I) was inhibited in five cases and with different degrees of inhibition. Moreover, the L. pneumophila sg. 1 used in this second screening was not susceptible to the activity of the antagonistic bacteria in four experiments, although all the antagonistic bacteria were originally selected for their ability to inhibit the growth of a reference strain of the same serogroup (L. pneumophila sg. 1 DSM7513). Finally, L. longbeachae and L. feeleii were the least susceptible strains, inhibited only by four antagonistic strains, of which three were the P. lurida isolates, while one was in one case P. iridis (L. feeleii) and in the other B. auriantiaca (II) (L. longbeachae).

### 3.2. Genome mining for potential inhibitory compounds

The genomes of our isolates were analysed with the software antiSMASH and revealed a broad range of Biosynthetic Gene Clusters (BGCs), of which Non-Ribosomal-Peptides (NRP) represented the main class. Table 2 reports the BGCs that showed high similarity (>60%) to previously known BGCs annotated in the antiSMASH database (22). The data shows how the three isolates of P. lurida that inhibited all the Legionella strains possessed a BGC with high similarity to the cluster encoding for the biosurfactant viscosin. Although the percent of similarity reported was different (100%; 75%; 68%), in two latter cases the BGC was located at the edge of a contig, not allowing for the full coverage of the cluster, and the missing parts were detected on other contigs for the two organisms.

Moreover, one strain of P. lurida (III) had three additional BGCs showing high similarity to flexirubin, rhizomide and icosalide respectively, which are to be attributed to the genome of the contaminant species. In fact, an analysis of the source microorganisms for these BGCs, conducted using the antiSMASH database, revealed their compatibility with the genus Chryseobacterium, which is the putative contaminant of the P. lurida (III) strain (see above).

The genome of P. iridis showed the presence of a BGC assigned to hydrogen cyanide, which is a volatile compound produced by some microorganisms with high toxicity towards other prokaryotic and eukaryotic organisms (23). While the production of a volatile compound would be able to explain the total inhibition observed in four cases with respect to P. iridis, it is notable that the same BGC was detected in the genome of P. lundensis, for which such total inhibition of Legionella was never observed. A possible explanation is that the BGC is only expressed by one strain, but no expression data is available for this work. The two strains of B. auriantiaca, as well as C. pauculus, did not reveal the presence of any high-similarity BGCs, highlighting the possibility of the inhibition being caused by a novel or orphan compound unassigned to any BGCs. BGCs for rhizomide and kolossin were detected in the genome of Pseudomonas_E sp900583165, while the cluster for zeaxanthin was present in the genome of Sphingomonas spp. While rhizomide is a cyclic-lipodepsipeptide (24), kolossin is a large non-ribosomal-peptide to our knowledge only isolated from the microorganism Photorhabdus luminescens (25). Zeaxanthin is a common carotenoid produced by plants and some microorganisms (26).

Subsequent work in our study focused on viscosin-producing bacteria, as the cluster was observed in multiple strains able to inhibit all the Legionella species tested and refers to a clear, characterized compound.

### 3.3. Identification of compounds belonging to the viscosin family in the cultures

In order to test whether the compounds causing the inhibition were secreted extracellularly, selected strains of Legionella (L. pneumophila sg.1; L. jordanis) were co-cultured with two strains of P. lurida (I; II), and the supernatant was tested against the same Legionella strains. L. jordanis was inhibited by the supernatant of all the combinations prepared, while no inhibition of L. pneumophila sg. 1 was observed, probably due to lower concentrations of the antagonistic compounds in the supernatant. Supernatants from co-cultures of L. jordanis and P. lurida (I), along with monocultures of Legionella and the antagonistic strain, were then subjected to liquid-liquid extraction followed by LC-MS analysis. This analysis identified masses [M+H]+ corresponding to viscosin (Retention Time [RT]: 19.33 min; 1126.69 m/z), with variable relative abundances ranging from 2E6 to 1.19E7 across all samples. Additionally, masses corresponding to the masses of Massetolide E (RT: 18.84 min; 1112.68 m/z; relative abundances 8.9E4 to 2.75E6) and Massetolide D (RT: 19.99 min; 1140.71 m/z; relative abundances 1.2E4 to 1.75E5) were detected. Complete LC-MS data details are provided in Supplementary Fig. S2-S4. MS/MS analysis further confirmed the molecular structure of viscosin in all samples where it was detected, as shown in Supplementary Fig. S2-S4

The supernatant of the co-culture L. jordanis – P. lurida (I) was afterwards analysed through preparative HPLC in order to separate the compounds and screen the fractions collected against all the Legionella species through spot-on-lawn assays. The results showed four fractions with antagonistic activity against all the Legionella strains (fractions 10, 14, 24, 25; Fig. 3). However, given that the eluate was collected at one-minute intervals rather than being synchronized with specific chromatographic peaks, it is conceivable that active compounds were distributed across multiple fractions, thereby manifesting inhibitory effects in consecutive fractions.

**Figure 3.**
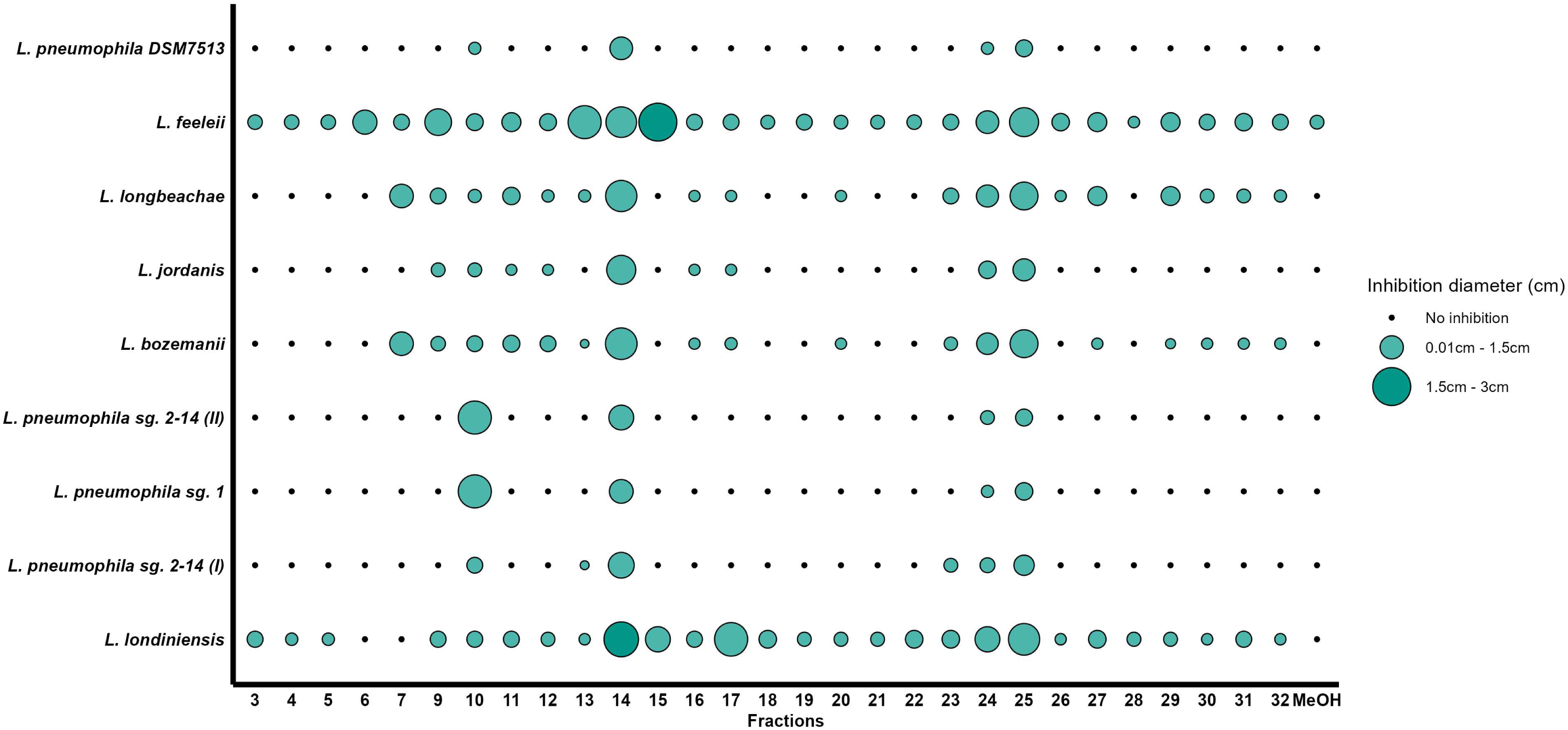
Results of the spot-on-lawn experiments in which the fractions obtained from the extracted supernatant, separated through HPLC (displayed in x-axis) were tested against the Legionella strains used in this work (displayed in y-axis). A green circle indicates that an inhibition was observed on the plate, while a black point is displayed when no inhibition was detected. All the experiments were performed in triplicate using blank antibiotic-susceptibility discs that were soaked with the respective fractions. The size of the green circles is proportional to the degree of the inhibition measured as the average inhibition zone diameter, normalized for the colony size

LC-MS analysis of the inhibitory fractions identified the presence masses [M+H]+ corresponding to viscosin in all four fractions (RT: 13 min; 1126.69 m/z; relative abundances 2E6 to 1.1E7). Masses corresponding to Massetolide E were detected in fractions 10 (RT: 12.40 min; 1112.68 m/z; relative abundance: 1.2E5) and fraction 25 (RT: 12.43 min; relative abundance: 5E3). Finally, a mass corresponding to the mass of Massetolide D were detected only in fraction 10 (RT: 12.63 min; 1140.71 m/z; relative abundance: 4.5E3). MS/MS confirmed the molecular structure of viscosin (Supplementary Fig. S5-S7). Notably, masses which could not be assigned to specific compounds were detected in fractions 10 and 14 (RT: 10.77 min; 460.36 m/z, relative abundances: 1.1E8-1.6E8) (RT: 13.23-13.35; 474.38 m/z; relative abundances: 1.7E7-1.35E8). An additional non-assigned mass was detected only in fraction 25 (RT: 7.66 min; 454 m/z; relative abundance 4.5E7).

## 4. Discussion

This study aimed at isolating bacteria with antagonistic properties against Legionella. We isolated 10 bacteria (predominantly belonging to the genus Pseudomonas) with antagonistic activity towards one reference strain of L. pneumophila, and observed variability in the inhibition pattern when we extended the screening to multiple Legionella species (Figs. 1, 2). Furthermore, we investigated the genomes of the antagonists to identify potential inhibitory compounds (Table 3) and were able to fractionate and verify the antimicrobial activity of the supernatant of a co-culture Legionella – P. lurida (Fig. 3), verifying the presence of the biosurfactant viscosin as the likely Legionella-antagonistic compound.

**Table 3.**
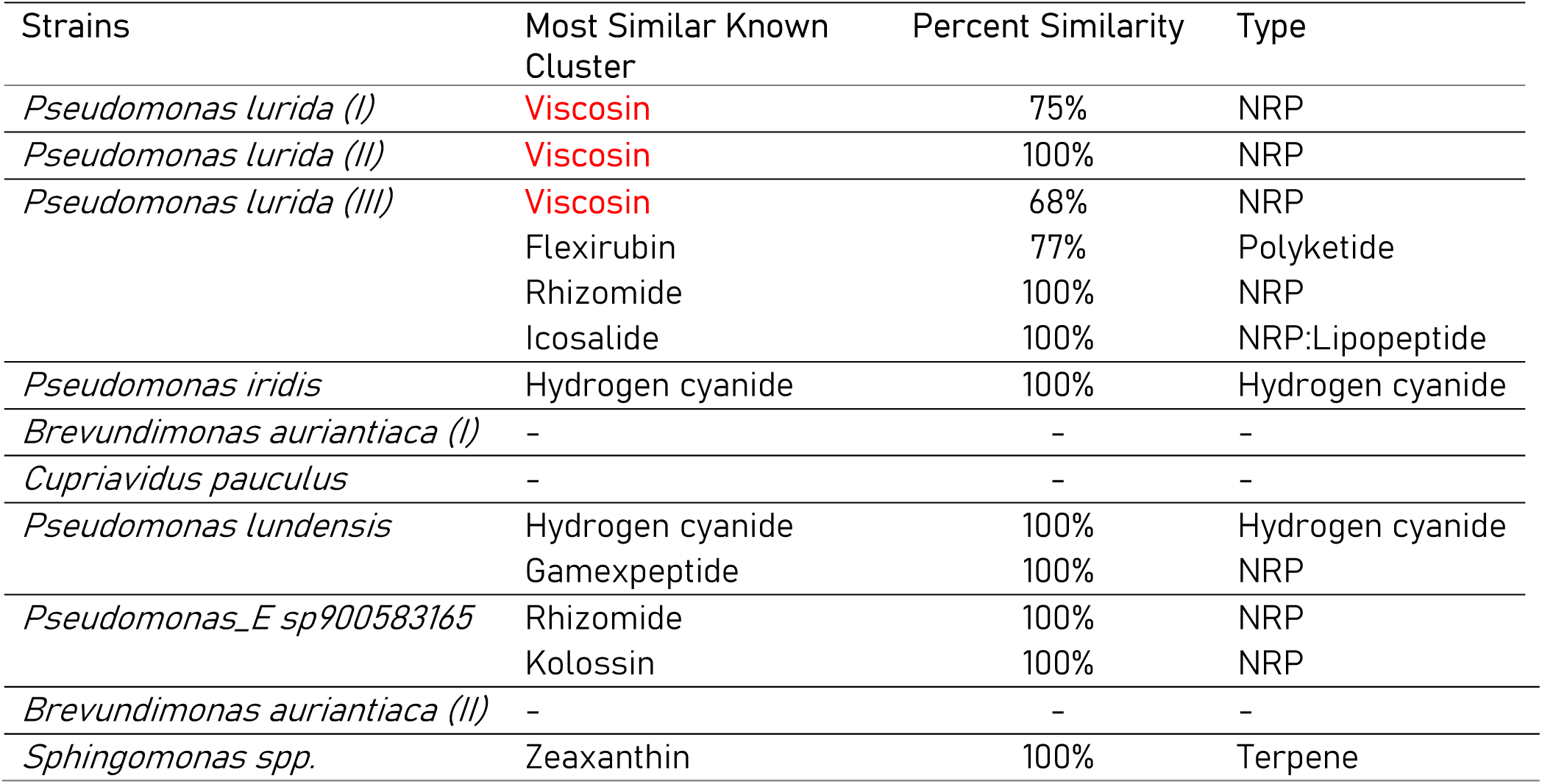
BGCs detected in the genomes of the antagonistic strains using antiSMASH. The BGCs have been selected based on the similarity to previously annotated clusters (>60% similarity).

### 4.1. Inhibition of Legionella spp. by aquatic bacteria

The study of microbial competition is an important tool for understanding how bacteria behave in stressful conditions, for the discovery of new antimicrobials and for elucidating underlying ecological mechanisms driving the establishment of microorganisms in specific environments (27, 28). In recent years, increased attention has been given to the competitive interactions between aquatic bacteria and Legionella, possibly due to the necessity to gain deeper ecological insights on a problematic opportunistic pathogen, and to investigate alternative mitigation strategies of this highly resistant genus (11, 29). Corre and coworkers (13) showed the antagonistic activity towards one strain of L. pneumophila by bacteria isolated from five different water sources, with a prevalence of antagonists belonging to the species Pseudomonas (n=70) and Aeromonas (n=19). Similar results were also presented by Guerrieri and coworkers (12), who showed the inhibition of several L. pneumophila strains by 80 aquatic bacteria, of which Pseudomonas represented the biggest fraction. Experiments performed using bacteria isolated from cooling towers described Bacillus spp., Chryseobacterium spp., Cupriavidus spp., Staphylococcus spp., and Stenotrophomonas spp. as antagonists of nine L. pneumophila strains (14).

We purposefully built on these studies, and isolated some bacterial species antagonistic towards Legionella from different water sources in Switzerland. We detected bacteria belonging to the genus Pseudomonas as the main Legionella antagonists (Fig. 2). An isolation bias is present, as many other aquatic bacteria (and potential antagonists) cannot grow in standard media and temperatures used in this and previous studies (13) . However, it is noticeable that previous molecular observations have also reported negative associations between Pseudomonas and Legionella in building plumbing systems (30, 31), cooling towers (32) and swimming pools (33). It is therefore possible that microorganisms belonging to these two genera share similar environmental preferences and occupy the same ecological niches, competing for available resources. Specifically related to the genus Pseudomonas, we report for the first time the antagonistic activity of the species P. lurida, P. iridis and P. lundensis, as well as Legionella inhibition by a novel Pseudomonas_E sp900583165. Combined with the data generated in other studies, this suggests that multiple Pseudomonas species can inhibit Legionella. Moreover, our experiments show anti-Legionella activity of bacteria belonging to the genus Brevundimonas, which have previously been associated with positive effects on the growth of L. pneumophila (34). These results suggest a variable outcome in the interactions between two genera, which can depend on specific species within the same genus (as our results also show in Fig. 2; discussed below), and on the general experimental and environmental conditions. Therefore, associations described by observational molecular studies, especially when the taxonomy is not resolved at the species level, might not reflect all the possible outcomes that interactions among members of different genera can have (35)

Overall, the results reported in this study provide additional new information about bacteria antagonistic to Legionella. However, the initial selection strategy for the antagonists was based on the inhibition of a reference strain of L. pneumophila. By doing this, given the variable inhibition reported in Fig. 2 (see Section 4.2), it is conceivable that we did not select for more potential antagonists towards other Legionella species, but also strains of L. pneumophila. This, together with the cultivation biases discussed above, calls for the use of improved isolation methods to identify a broader range of antagonists. In this context, pairing molecular observations with laboratory experiments represents a valuable strategy. Through molecular-based work, it is possible to obtain an identification of the microorganisms correlating with the absence of Legionella (31, 34, 35), and then try to isolate those or use cultures from strain collections. However, due to the limited taxonomic information often provided by these approaches (especially for 16S amplicon sequencing), the isolation of a specific strain remains challenging. Here we demonstrate that whole genome sequencing provides information regarding the specific functions associated with Legionella antagonism, helping the understanding what represents an unfavorable niche for these bacteria. Moreover, our study provide access to the gene clusters, which can be heterologously expressed, into aquatic microorganisms easy to manipulate and verify their biological activity: this would overcome the problems related to the isolation of the antagonistic strains.

### 4.2. Variability in the inhibition patterns highlights **Legionella** diversity and potential ecological characteristics

In the environment, bacteria compete for space and resources, using either interference or exploitative competition (36). The results of these competitive interactions are likely to have a strong influence on the composition of communities and how these are distributed in the environment (28, 37). We showed that Legionella species are inhibited by a range of antagonistic bacteria, but also that the inhibition presented inter-species and intra-species variability (Fig. 2). This variability in the inhibition pattern can be analysed from an ecological perspective, providing some useful insights. For instance, Russel and colleagues (38) demonstrated that bacteria are likely to exhibit antagonism towards microorganisms that present metabolic similarities, hence sharing nutritional preferences. In the context of our study, given the already discussed negative relationship between Pseudomonas and Legionella, this would reinforce the hypothesis that the two genera share similar niches and probably share some nutritional requirements. This would especially be relevant for the extracellular phase of the lifecycle of Legionella, given especially its preference for dense environments likely occupied by Pseudomonas such as biofilms (3). Moreover, this interpretation of the obtained results suggests the possibility that different Legionella species and strains live within distinct ecological niches in the environment. In a previous published study from our group (35), we have already observed a high genetic variability in the 16S amplicon sequence variants associated with the genus Legionella, as well as distinct patterns in the way the various Legionella ASVs correlated amongst themselves and the rest of the community, which would once more support the idea that different ecological niches are occupied. This would explain co-occurrence within the same larger environment. These observations would be in agreement with multiple studies suggesting that differences between related groups of bacteria are driven by the way they differentiate in the environment (5, 39–41).

Substantial information exists regarding the antagonistic capabilities of Pseudomonas, which is a genus of bacteria known to harbour many different inhibition strategies (15, 42). In this context, it has already been demonstrated that Pseudomonas strains isolated from different environmental niches show variable antagonistic activity (43). Notably, even at the species level, significant differences in the outcome of competitive interactions are present, as shown by Lyng and Kovàcs (44) when assessing the interactions between Pseudomonas and Bacillus. The same authors described broad diversity in secondary metabolites responsible for antagonism across members of the same species, and even more of the same genus, hypothesising that differences in inhibition might be driven by acquisition of genes through horizontal gene transfer. Microorganisms have also evolved multiple traits to defend against the attack of their neighbours (45) and this translates to the possibility that variable outcomes in competition are not only due to different compounds produced by the antagonists. For example, in our study, we observed that some antagonists could inhibit some Legionella, but not others (Fig. 2); at the same time, in multiple cases one Legionella species is inhibited by several different antagonists. The first situation suggests that Legionella species can use various resistance mechanisms, while the multitude of compounds produced by the antagonists can be responsible for the second scenario.

Regardless of the exact reason behind the variable outcome of the competitive interactions between Legionella and the antagonists found in this and other studies (whether related to the environment or specific antagonistic/defence mechanisms), differences are present at the species and strain level for the genus Legionella. In a recent study (5), we conducted a genome-wide investigation of the family Legionellaceae, and reported a high level of diversity related to number of species, cellular components and metabolism. While this data indicates different behaviours in the environment, to our knowledge, genus-level differences of the responses towards antagonistic mechanisms have not been explored in a broader ecological perspective for Legionella, although this is important because of the pathogenic nature of many members of the genus (4). Purposefully, the approach undertaken in this study aimed at exploring the response of multiple Legionella species to antagonistic bacteria, bringing the experiment beyond the only use of L. pneumophila. This type of ecological information will support distinguishing between species, and provide insights on pathogenicity, control strategies and sampling/detection approaches.

A few studies have also started to predict broader ecological information on community assembly from pairwise inhibition building models (46) and dedicated laboratory experiments (47). While our study was not designed to allow that such predictions could be extrapolated (specifically for the lack of quantitative growth data), collecting the necessary information in order to generate these predictions represents an important step for future experiments. It could provide valuable information on the microbial interactions and exclusive relationships between Legionella and other microorganisms in engineered aquatic environments, which could be used in turn to design alternative mitigation strategies for Legionella based on the use of live competitors (29). As for all the studies that deal with competition, considerations about higher-order-interactions (48) must be made, since many of the ecological theories are built using pairwise interactions. Moreover, all these microbial interactions should be analysed in the light of the intracellular life cycle that Legionella go through in the environment inside protists (3), which can give a different meaning to these competition experiments.

### 4.3. The role of biosurfactants in the inhibition of **Legionella**

Bacterial competition, and in particular interference competition, is often manifested through the secretion of several antibacterial compounds (49). A few studies investigated the molecules responsible for the inhibition. Faucher and coworkers showed that a strain of P. alcaliphila, commonly found in cooling towers, was able to inhibit Legionella through the production of toxoflavin (50). Corre and coworkers identified the anti-Legionella volatile compound 1-undecene (51), while the inhibitory activity of the novel compound warnericin from Staphylococcus warnerii was first described by Héchard and coworkers (52). In two subsequent studies, Loiseau and coworkers first showed that surfactin was able to inhibit multiple Legionella species which presented a low minimum inhibitory concentration compared to other gram-positive and -negative bacteria (53), and then illustrated the strong activity of multiple biosurfactants (e.g. rhamnolipids and lipopeptides mixtures) produced by Pseudomonas strains (16).

Consistently with these findings, we detected BGCs related to the biosurfactants viscosin, rhizomide and icosalide in the genomes of the antagonists tested (Table 2). Interestingly, the three strains of P. lurida that exhibited the broadest spectrum of inhibition towards the Legionella species tested (Fig. 2) were the only ones to possess the cluster for viscosin. The fact that we detected the masses corresponding to the masses of viscosin and massetolides A and D (belonging to the same viscosin family) in the supernatant and in the fractions obtained through HPLC, is therefore a strong indication of the fact that viscosin is indeed the compound responsible for the inhibition observed. Viscosin is a lipopeptide produced by several Pseudomonas spp. by a non-ribosomal peptide synthetase with demonstrated antimicrobial and antiviral activity (54), and is involved in the processes of biofilm formation and dispersal (55). Interestingly, the mode of action of several anti-Legionella compounds (including biosurfactants) is directed towards the bacterial membrane, suggesting a high susceptibility of Legionella to these molecules (11, 16). Moreover, while biosurfactants seem to be mostly active towards gram-positive bacteria (56), Legionella represents an exception in this sense highlighting that the composition of their membrane may drive increased susceptibility.

Biosurfactants have recently gained attention due to their industrial potential, and have been increasingly used in food, cosmetic, pharmaceutical and oil/gas remediation fields as an alternative to chemical surfactants, which present higher toxicity and environmental impact (57). In the environment, biosurfactants are produced by certain genera of microorganisms for reasons related to motility, competition and especially biofilm formation/removal (58). Therefore, it is conceivable, given the biofilm-associated nature of Legionella (10), that the pathogen enters in contact with these compounds in the environment, and that their effects are not only limited to what is observed in laboratory experiments.

However, it is not possible to exclude that the inhibition of Legionella was caused by other compounds (alone or in combination with biosurfactants). Our genomic data suggest the presence of multiple BGCs (Table 3), hence, several molecules might be responsible for the inhibition observed, especially in those cases of variable Legionella inhibition. While we did not search for additional compounds in this study, the opportunity exists to conduct more targeted experiments in order to identify more anti-Legionella molecules to add to the list that many above-mentioned studies have contributed, and to ultimately test the biological activity in more realistic settings with dedicated laboratory or pilot scale experiments.

Given the importance of reducing the levels of Legionella in engineered aquatic ecosystems, the use of new compounds will represent an alternative to chemicals (e.g., chlorine) whose efficacy against Legionella is lacking consistency. The adoption of new compounds with low toxicity might also help the acceptance of water disinfection in countries (e.g., Switzerland or the Netherlands) where this is less common. However, a few considerations are necessary in this regard. Compounds of biological origin, although in many cases less toxic than chemically synthetized ones, are only obtained in small amounts due to their production process (58) and are therefore currently not affordable by the water industries. Moreover, public acceptance for such alternatives is likely difficult to obtain, and these approaches would rather find application in systems working with water that will not directly enter in contact with the public (e.g., cooling towers, process water). Finally, these mitigation strategies would not be effective against Legionella when they are internalized in their eukaryotic hosts. Nevertheless, some of these compounds (as demonstrated for biosurfactants) can promote the disruption of the biofilms (55, 58), overcoming the challenges related to the intracellular Legionella.

## 5. Conclusions

- Multiple bacteria were able to inhibit different species and strains of Legionella, but the interaction results were variable in both outcome and degree of inhibition, demonstrating the importance of considering Legionella species diversity in antagonism studies.
- Four species of Pseudomonas were described as Legionella antagonists for the first time in this study.
- Variable inhibition of Legionella by other strains and compounds could explain important ecological differences among Legionella species in the environment. This is relevant given the occurrence of multiple pathogenic Legionella species in the environment. However, dedicated additional studies are necessary to reinforce this hypothesis.
- The biosurfactant viscosin was identified as the compounds potentially responsible for the broad inhibition of Legionella, supporting previous observations that biosurfactants are interesting anti-Legionella compounds.

## Acknowledgements

This research was funded by the Federal Food Safety and Veterinary Office (FSVO), in partnership with the Federal Offices of Public Health (FOPH) and Energy (SFOE) in Switzerland, through the project LeCo (Legionella Control in Buildings; Aramis nr.:4.20.01) and Eawag discretionary funding. Alessio Cavallaro acknowledges Innosuisse for support through the project PROMOTE (Project nr.: 115.167.1 IP-EE. Silke Probst acknowledges the Helmut Horten Foundation for support (project ID: 2022-YIG-090).

## References

1. Fields BS, Benson RF, Besser RE. 2002. Legionella and Legionnaires’ disease: 25 years of investigation. Clin Microbiol Rev 15:506–26.

2. Falkinham JO, 3rd, Hilborn ED, Arduino MJ, Pruden A, Edwards MA. 2015. Epidemiology and Ecology of Opportunistic Premise Plumbing Pathogens: Legionella pneumophila, Mycobacterium avium, and Pseudomonas aeruginosa. Environ Health Perspect 123:749–58.

3. Mondino S, Schmidt S, Rolando M, Escoll P, Gomez-Valero L, Buchrieser C. 2020. Legionnaires’ Disease: State of the Art Knowledge of Pathogenesis Mechanisms of Legionella. Annu Rev Pathol 15:439–466.

4. Chambers ST, Slow S, Scott-Thomas A, Murdoch DR. 2021. Legionellosis Caused by Non- Legionella pneumophila Species, with a Focus on Legionella longbeachae. Microorganisms 9.

5. Gabrielli M, Cavallaro A, Hammes F. 2024. Expansion, restructuring and characterization of the *Legionellaceae* family. bioRxiv doi:10.1101/2024.10.21.619444:2024.10.21.619444.

6. Muder RR, Victor LY. 2002. Infection due to Legionella species other Than L. pneumophila. Clinical Infection Diseases 35:990–998.

7. Chauhan D, Shames SR. 2021. Pathogenicity and Virulence of Legionella: Intracellular replication and host response. Virulence 12:1122–1144.

8. Dilger T, Melzl H, Gessner A. 2018. Legionella contamination in warm water systems: A species-level survey. Int J Hyg Environ Health 221:199–210.

9. Nhu Nguyen TM, Ilef D, Jarraud S, Rouil L, Campese C, Che D, Haeghebaert S, Ganiayre F, Marcel F, Etienne J, Desenclos J-C. 2006. A Community-Wide Outbreak of Legionnaires Disease Linked to Industrial Cooling Towers—How Far Can Contaminated Aerosols Spread? The Journal of Infectious Diseases 193:102–111.

10. Declerck P. 2010. Biofilms: the environmental playground of Legionella pneumophila. Environ Microbiol 12:557–66.

11. Berjeaud JM, Chevalier S, Schlusselhuber M, Portier E, Loiseau C, Aucher W, Lesouhaitier O, Verdon J. 2016. Legionella pneumophila: The Paradox of a Highly Sensitive Opportunistic Waterborne Pathogen Able to Persist in the Environment. Front Microbiol 7:486.

12. Guerrieri E, Bondi M, Sabia C, de Niederhausern S, Borella P, Messi P. 2008. Effect of bacterial interference on biofilm development by Legionella pneumophila. Curr Microbiol 57:532–6.

13. Corre MH, Delafont V, Legrand A, Berjeaud JM, Verdon J. 2018. Exploiting the Richness of Environmental Waterborne Bacterial Species to Find Natural Legionella pneumophila Competitors. Front Microbiol 9:3360.

14. Paranjape K, Levesque S, Faucher SP. 2022. Bacterial Antagonistic Species of the Pathogenic Genus Legionella Isolated from Cooling Tower. Microorganisms 10.

15. Gross H, Loper JE. 2009. Genomics of secondary metabolite production by Pseudomonas spp. Nat Prod Rep 26:1408–46.

16. Loiseau C, Portier E, Corre MH, Schlusselhuber M, Depayras S, Berjeaud JM, Verdon J. 2018. Highlighting the Potency of Biosurfactants Produced by Pseudomonas Strains as Anti- Legionella Agents. Biomed Res Int 2018:8194368.

17. Prjibelski A, Antipov D, Meleshko D, Lapidus A, Korobeynikov A. 2020. Using SPAdes De Novo Assembler. Current Protocols in Bioinformatics 70:e102.

18. Chklovski A, Parks DH, Woodcroft BJ, Tyson GW. 2023. CheckM2: a rapid, scalable and accurate tool for assessing microbial genome quality using machine learning. Nature Methods 20:1203–1212.

19. Orakov A, Fullam A, Coelho LP, Khedkar S, Szklarczyk D, Mende DR, Schmidt TSB, Bork P. 2021. GUNC: detection of chimerism and contamination in prokaryotic genomes. Genome Biology 22:178.

20. Chaumeil P-A, Mussig AJ, Hugenholtz P, Parks DH. 2022. GTDB-Tk v2: memory friendly classification with the genome taxonomy database. Bioinformatics 38:5315–5316.

21. Blin K, Shaw S, Augustijn HE, Reitz ZL, Biermann F, Alanjary M, Fetter A, Terlouw BR, Metcalf WW, Helfrich EJN, van Wezel GP, Medema MH, Weber T. 2023. antiSMASH 7.0: new and improved predictions for detection, regulation, chemical structures and visualisation. Nucleic Acids Research 51:W46–W50.

22. Blin K, Shaw S, Medema MH, Weber T. 2023. The antiSMASH database version 4: additional genomes and BGCs, new sequence-based searches and more. Nucleic Acids Research 52:D586–D589.

23. Blumer C, Haas D. 2000. Mechanism, regulation, and ecological role of bacterial cyanide biosynthesis. Arch Microbiol 173:170–7.

24. Wang X, Zhou H, Chen H, Jing X, Zheng W, Li R, Sun T, Liu J, Fu J, Huo L, Li Y-z, Shen Y, Ding X, Müller R, Bian X, Zhang Y. 2018. Discovery of recombinases enables genome mining of cryptic biosynthetic gene clusters in Burkholderiales species. Proceedings of the National Academy of Sciences 115:E4255–E4263.

25. Bode HB, Brachmann AO, Jadhav KB, Seyfarth L, Dauth C, Fuchs SW, Kaiser M, Waterfield NR, Sack H, Heinemann SH, Arndt H-D. 2015. Structure Elucidation and Activity of Kolossin A, the D-/L-Pentadecapeptide Product of a Giant Nonribosomal Peptide Synthetase. Angewandte Chemie International Edition 54:10352–10355.

26. Sajilata MG, Singhal RS, Kamat MY. 2008. The Carotenoid Pigment Zeaxanthin—A Review. Comprehensive Reviews in Food Science and Food Safety 7:29–49.

27. Mullis MM, Rambo IM, Baker BJ, Reese BK. 2019. Diversity, Ecology, and Prevalence of Antimicrobials in Nature. Front Microbiol 10:2518.

28. Ghoul M, Mitri S. 2016. The Ecology and Evolution of Microbial Competition. Trends Microbiol 24:833–845.

29. Cavallaro A, Rhoads WJ, Huwiler SG, Stachler E, Hammes F. 2022. Potential probiotic approaches to control Legionella in engineered aquatic ecosystems. FEMS Microbiol Ecol 98.

30. Proctor CR, Reimann M, Vriens B, Hammes F. 2018. Biofilms in shower hoses. Water Res 131:274–286.

31. Scaturro M, Chierico FD, Motro Y, Chaldoupi A, Flountzi A, Moran-Gilad J, Girolamo A, Koutsiomani T, Krogulska B, Lindsay D, Matuszewska R, Papageorgiou G, Pancer K, Panoussis N, Rota MC, Uldum SA, Velonakis E, Chaput DL, Ricci ML. 2022. Bacterial communities of premise plumbing systems in four European cities, and their association with culturable Legionella. bioRxiv doi:10.1101/2022.08.12.503735.

32. Paranjape K, Bedard E, Whyte LG, Ronholm J, Prevost M, Faucher SP. 2020. Presence of Legionella spp. in cooling towers: the role of microbial diversity, Pseudomonas, and continuous chlorine application. Water Res 169:115252.

33. Leoni E, Legnami PP, Bucci Sabattini MA, Righi F. 2001. Prevalence of Legionella spp. in swimming pool environment. Water Research 15:3749–3753.

34. Paranjape K, Bedard E, Shetty D, Hu M, Choon FCP, Prevost M, Faucher SP. 2020. Unravelling the importance of the eukaryotic and bacterial communities and their relationship with Legionella spp. ecology in cooling towers: a complex network. Microbiome 8:157.

35. Cavallaro A, Rhoads WJ, Sylvestre E, Marti T, Walser JC, Hammes F. 2023. Legionella relative abundance in shower hose biofilms is associated with specific microbiome members. FEMS Microbes 4:xtad016.

36. Granato ET, Meiller-Legrand TA, Foster KR. 2019. The Evolution and Ecology of Bacterial Warfare. Curr Biol 29:R521–R537.

37. Hibbing ME, Fuqua C, Parsek MR, Peterson SB. 2010. Bacterial competition: surviving and thriving in the microbial jungle. Nat Rev Microbiol 8:15–25.

38. Russel J, Roder HL, Madsen JS, Burmolle M, Sorensen SJ. 2017. Antagonism correlates with metabolic similarity in diverse bacteria. Proc Natl Acad Sci U S A 114:10684–10688.

39. Patin NV, Duncan KR, Dorrestein PC, Jensen PR. 2016. Competitive strategies differentiate closely related species of marine actinobacteria. ISME J 10:478–90.

40. Johnson ZI, Zinser ER, Coe A, McNulty NP, Woodward EMS, Chisholm SW. 2006. Niche Partitioning Among *Prochlorococcus* Ecotypes Along Ocean-Scale Environmental Gradients. Science 311:1737–1740.

41. Yawata Y, Cordero OX, Menolascina F, Hehemann J-H, Polz MF, Stocker R. 2014. Competition– dispersal tradeoff ecologically differentiates recently speciated marine bacterioplankton populations. Proceedings of the National Academy of Sciences 111:5622–5627.

42. Leinweber A, Weigert M, Kümmerli R. 2018. The bacteriumPseudomonas aeruginosasenses and gradually responds to interspecific competition for iron. Evolution 72:1515–1528.

43. Chiellini C, Lombardo K, Mocali S, Miceli E, Fani R. 2019. Pseudomonas strains isolated from different environmental niches exhibit different antagonistic ability. Ethology Ecology & Evolution 31:399–420.

44. Lyng M, Kovacs AT. 2023. Frenemies of the soil: Bacillus and Pseudomonas interspecies interactions. Trends Microbiol 31:845–857.

45. Zhou G, Shi QS, Huang XM, Xie XB. 2015. The Three Bacterial Lines of Defense against Antimicrobial Agents. Int J Mol Sci 16:21711–33.

46. Lee H, Bloxham B, Gore J. 2023. Resource competition can explain simplicity in microbial community assembly. Proc Natl Acad Sci U S A 120:e2212113120.

47. Schmitz DA, Wechsler T, Mignot I, Kümmerli R. 2024. Predicting bacterial interaction outcomes from monoculture growth and supernatant assays. ISME Communications doi:10.1093/ismeco/ycae045.

48. Bairey E, Kelsic ED, Kishony R. 2016. High-order species interactions shape ecosystem diversity. Nat Commun 7:12285.

49. Peterson SB, Bertolli SK, Mougous JD. 2020. The Central Role of Interbacterial Antagonism in Bacterial Life. Curr Biol 30:R1203–R1214.

50. Faucher SP, Matthews S, Nickzad A, Vounba P, Shetty D, Bédard É, Prévost M, Déziel E, Paranjape K. 2022. Toxoflavin secreted by Pseudomonas alcaliphila inhibits growth of Legionella pneumophila and its host Vermamoeba vermiformis. BioRxiv doi:10.1101/2022.01.08.475489.

51. Corre M-H, Mercier A, Bouteiller M, Khalil A, Ginevra C, Depayras S, Dupont C, Rouxel M, Gallique M, Grac L, Jarraud S, Giron D, Merieau A, Berjeaud J-M, Verdon J, Goldberg JB. 2021. Bacterial Long-Range Warfare: Aerial Killing of Legionella pneumophila by Pseudomonas fluorescens. Microbiology Spectrum 9.

52. Hechard Y, Ferraz S, Bruneteau E, Steinert M, Berjeaud JM. 2005. Isolation and characterization of a Staphylococcus warneri strain producing an anti-Legionella peptide. FEMS Microbiol Lett 252:19–23.

53. Loiseau C, Schlusselhuber M, Bigot R, Bertaux J, Berjeaud JM, Verdon J. 2015. Surfactin from Bacillus subtilis displays an unexpected anti-Legionella activity. Appl Microbiol Biotechnol 99:5083–93.

54. Neu TR, Härtner T, Poralla K. 1990. Surface active properties of viscosin: a peptidolipid antibiotic. Applied Microbiology and Biotechnology 32:518–520.

55. Bonnichsen L, Bygvraa Svenningsen N, Rybtke M, de Bruijn I, Raaijmakers JM, Tolker-Nielsen T, Nybroe O. 2015. Lipopeptide biosurfactant viscosin enhances dispersal of Pseudomonas fluorescens SBW25 biofilms. Microbiology (Reading) 161:2289–97.

56. Raaijmakers JM, De Bruijn I, Nybroe O, Ongena M. 2010. Natural functions of lipopeptides from Bacillus and Pseudomonas: more than surfactants and antibiotics. FEMS Microbiol Rev 34:1037–62.

57. Nikolova C, Gutierrez T. 2021. Biosurfactants and Their Applications in the Oil and Gas Industry: Current State of Knowledge and Future Perspectives. Front Bioeng Biotechnol 9:626639.

58. Sharma J, Sundar D, Srivastava P. 2021. Biosurfactants: Potential Agents for Controlling Cellular Communication, Motility, and Antagonism. Front Mol Biosci 8:727070.

