## Supplementary tables and figures for "Variable inhibition of different Legionella species by antagonistic bacteria"

**Table 1.** List of the sources and relative location for the water samples used in the present study. Every water sample was plated in two different growth media (LB medium and R2A medium). 1 mL of each sample was split into five different plates (200 μL per plate). The table also reports the number of isolates that showed inhibitory activity towards a reference strain of L. pneumophila DSM7513

| Source | Location | Nr. of inhibitory isolates |
| --- | --- | --- |
| Tap water | Zurich (CH) | 0 |
| Tap water | Basel (CH) | 0 |
| Tap water | Dübendorf (CH) | 2 |
| Shower water | Zurich (CH) | 4 |
| Shower water | Basel (CH) | 6 |
| Shower water | Dübendorf (CH) | 1 |
| Lake water | Zurich (CH) | 2 |
| Pond water | Dübendorf (CH) | 19 |
| Groundwater | Dübendorf (CH) | 0 |
| Bottled water | - | 0 |

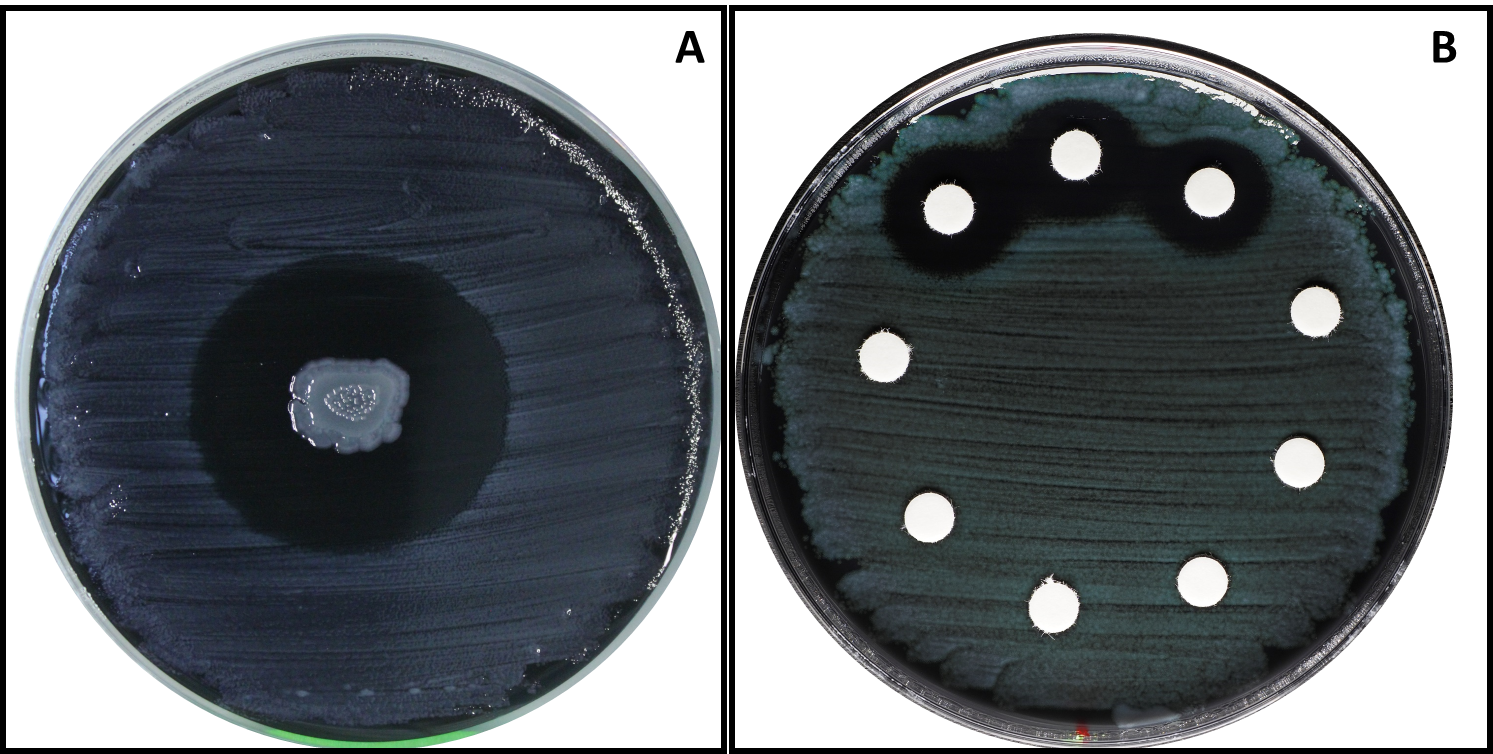

**Figure 1.** The figure shows examples of the two approaches used for the spot-on-lawn experiments performed in this study. A) An example of the spot-on-lawn conducted with the antagonistic bacteria. The antagonist (colony in the middle) is spotted on top of a lawn of Legionella; B) In order to test the fractions, blank antibiotic susceptibility discs were soaked with the solution of interest and, once dry, placed on a lawn of Legionella. In both cases, an inhibition is observed when Legionella does not grow around the colony/disc.

**Table 2.** Specific settings used for LC-MS analysis in all the experiments reported in this study. When applicable, values are given with their respective units.

| Setting | Value |
| --- | --- |
| Spray voltage | 3.5 kV |
| Capillary temperature | 320°C |
| Sheath gas | 57.5 |
| Aux gas | 16.25 |
| Spare gas | 3.25 |
| Probe heater | 462.50°C |
| Mode | Positive |
| Resolution | 30,000 |
| Microscans | 1 |
| Maximum IT | 100ms |
| Scan range | 150-200 m/z |

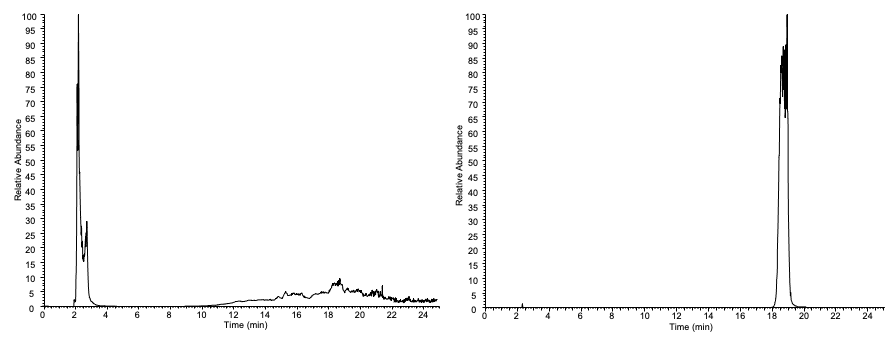

**A**

**B**

**C**

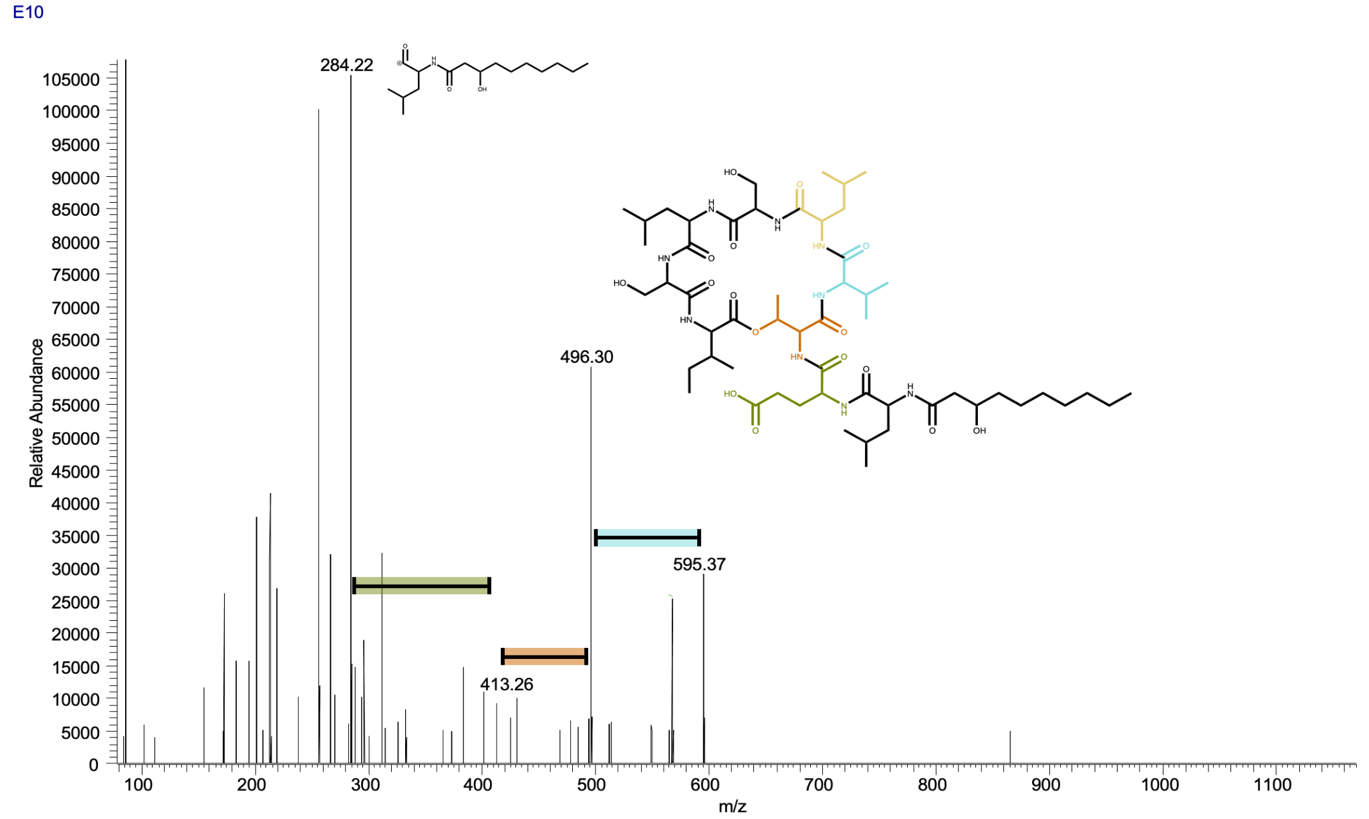

**Figure 2.** HR-LC-ESIMS data of the full *Pseudomonas lurida* (I) extract A) Total Ion Chromatogram (TIC) of the extracted culture. B) Extracted Ion Chromatogram (EIC) (m/z 1126.67 [M+H]) of viscosin. C) Measured fragments and ESI-MS/MS-spectrum of viscosin.

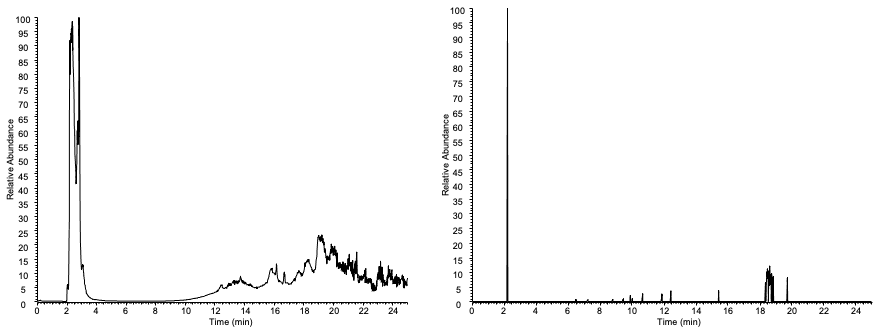

**B**

**A**

**Figure 3.** HR-LC-ESIMS data of the full *Legionella jordanis* extract A) Total Ion Chromatogram (TIC) of the extracted culture. B) Extracted Ion Chromatogram (EIC) (m/z 1126.67 [M+H]) of viscosin.

**B**

**A**

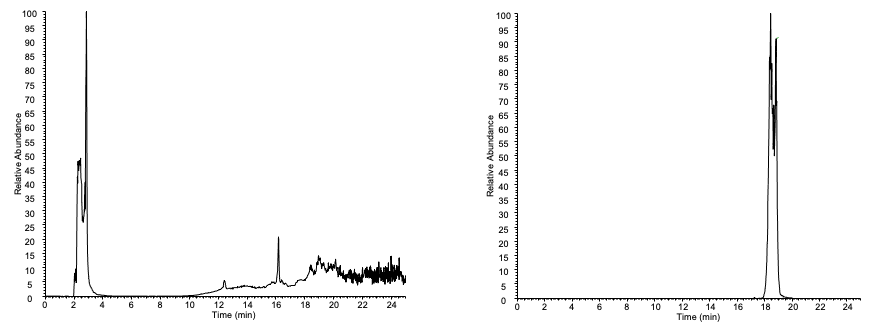

**C**

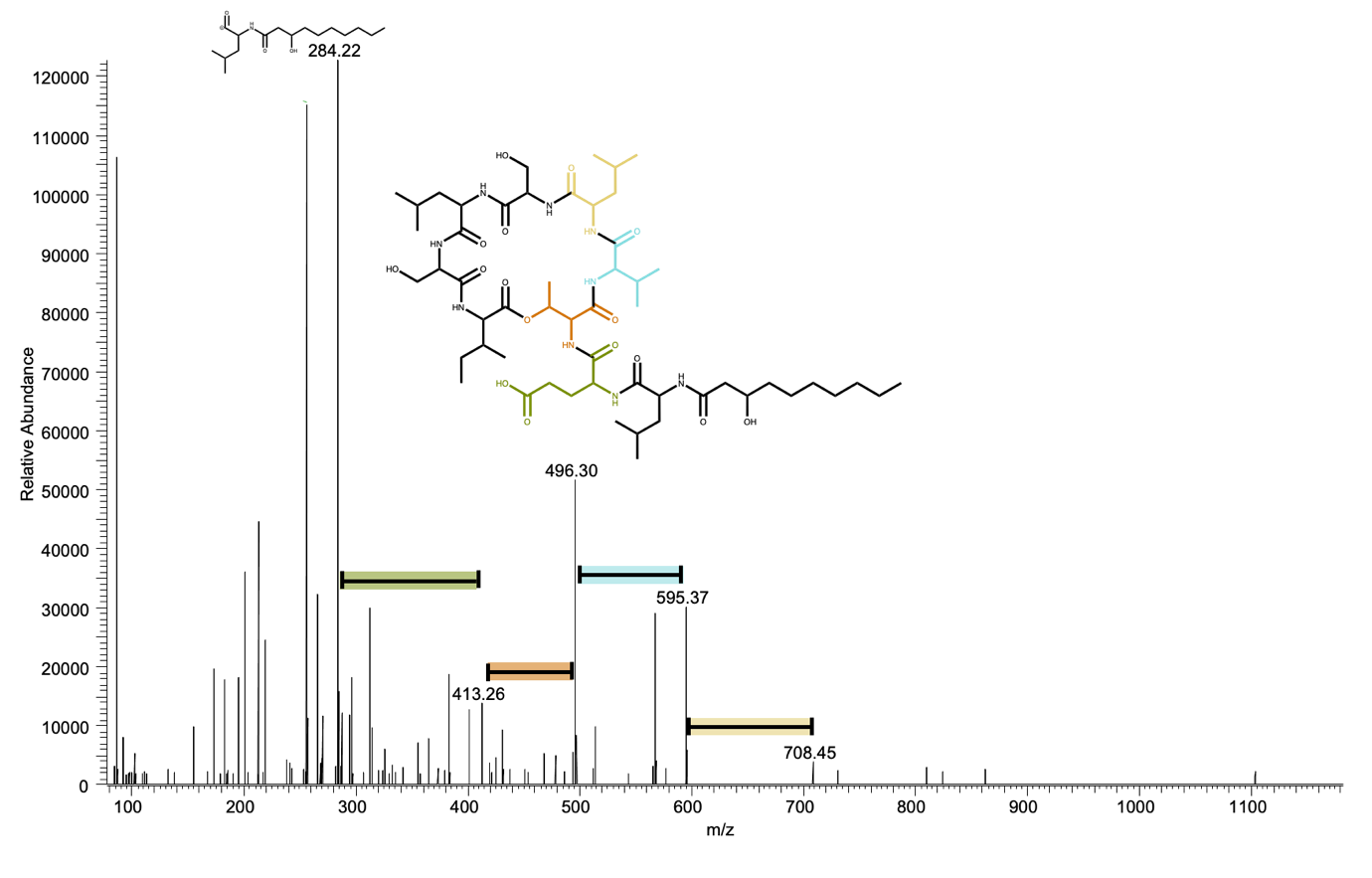

**Figure 4.** HR-LC-ESIMS data of the full *Pseudomonas* lurida (I) - *Legionella* jordanis co-culture extract A) Total Ion Chromatogram (TIC) of the extracted co-culture. B) Extracted Ion Chromatogram (EIC) (m/z 1126.67 [M+H]) of viscosin. C) Measured fragments and ESI-MS/MS-spectrum of viscosin.

**B**

**A**

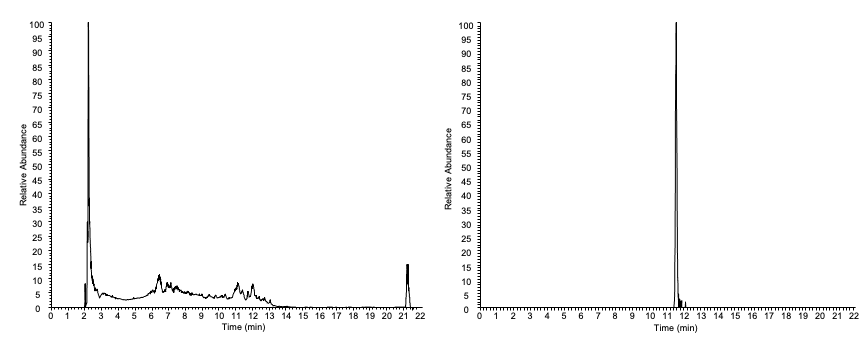

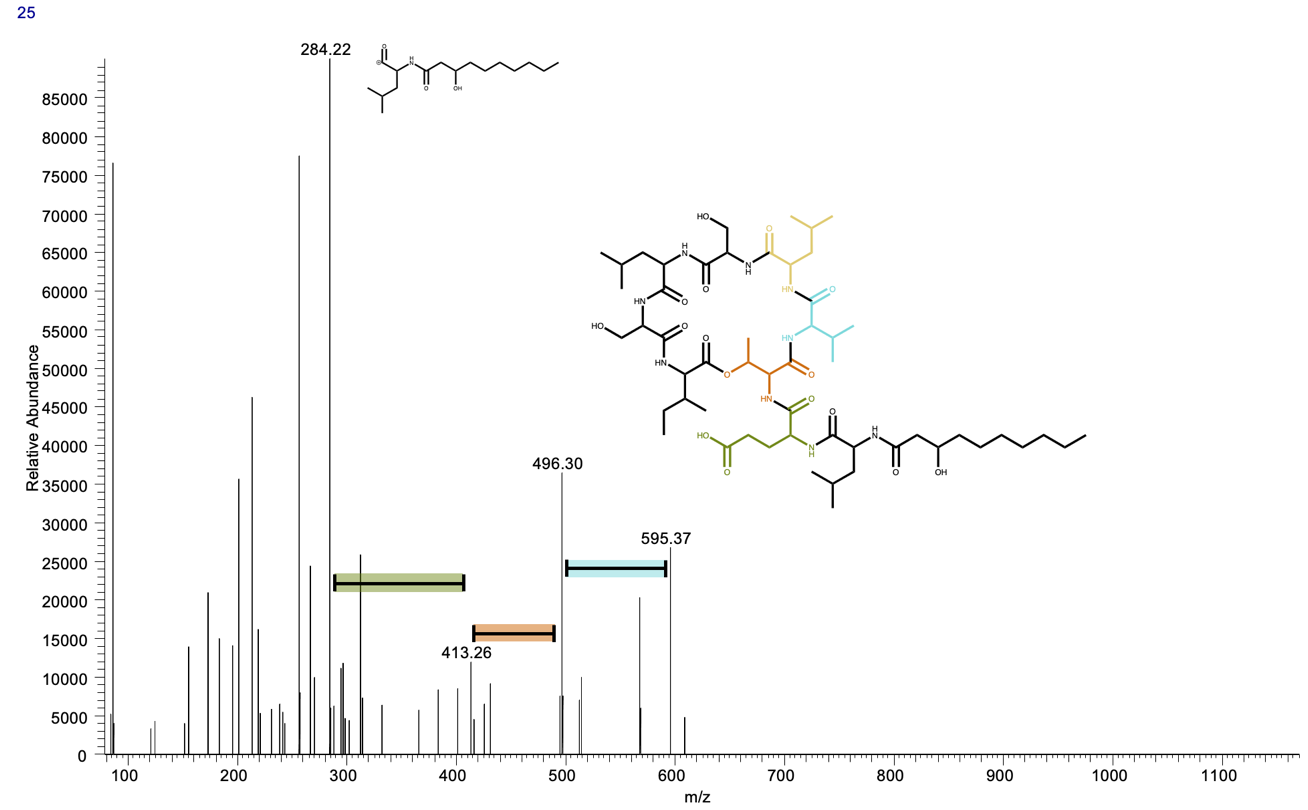

**C**

**Figure 5.** HR-LC-ESIMS data of the fraction 24 of the *Pseudomonas* lurida (I) -*Legionella* jordanis co-culture extract A) Total Ion Chromatogram (TIC) of the fraction 24 of the extracted co-culture (E10+18). B) Extracted Ion Chromatogram (EIC) (m/z 1126.67 [M+H]) of viscosin. C) Measured fragments and ESI-MS/MS-spectrum of viscosin.

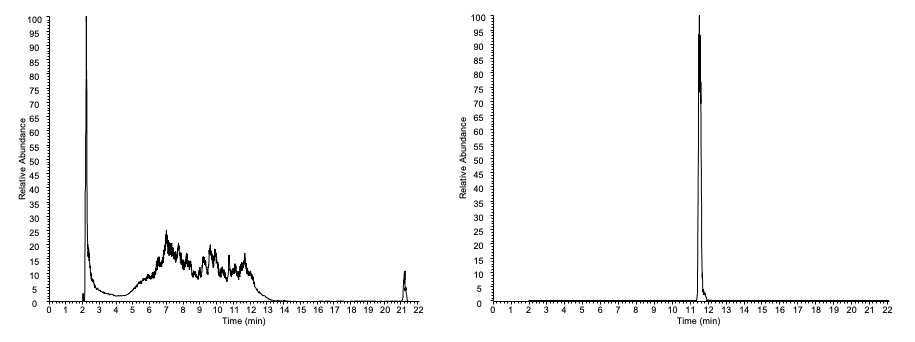

**B**

**A**

**C**

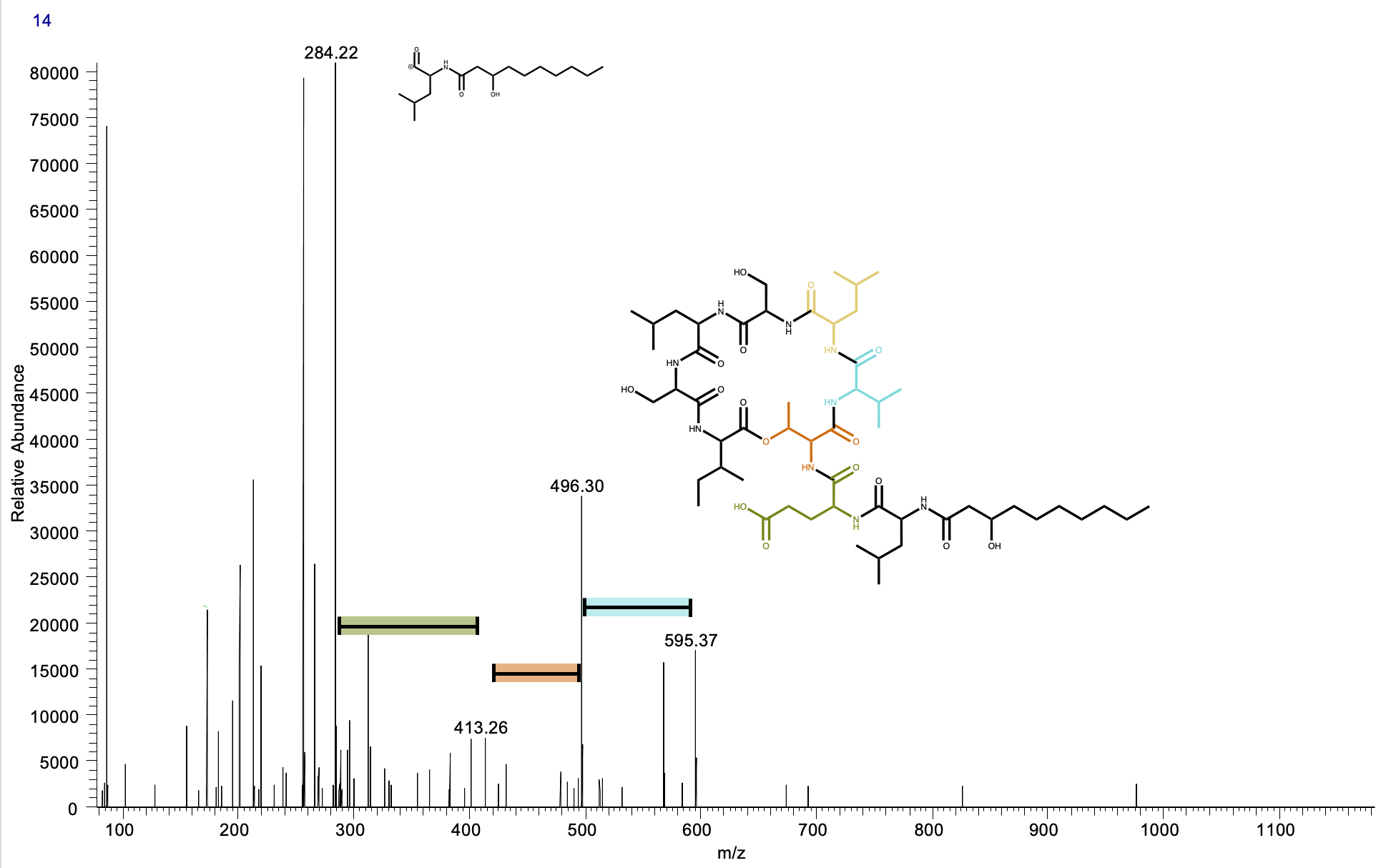

**Figure 6.** HR-LC-ESIMS data of the fraction 14 of the *Pseudomonas* *lurida* (I) - *Legionella* *jordanis* co-culture extract A) Total Ion Chromatogram (TIC) of the fraction 14 of the extracted co-culture. B) Extracted Ion Chromatogram (EIC) (m/z 1126.67 [M+H]) of viscosin. C) Measured fragments and ESI-MS/MS-spectrum of viscosin.

**B**

**A**

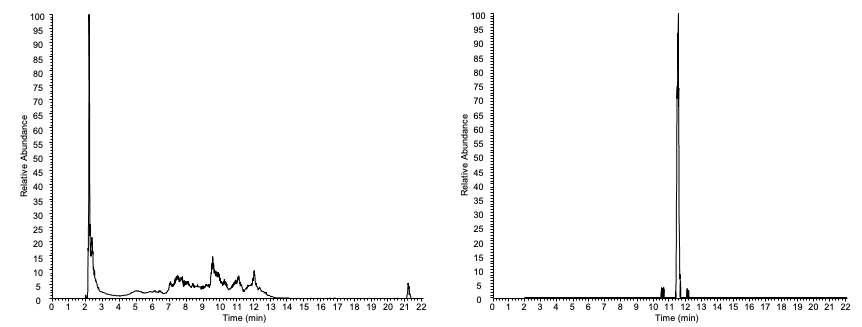

**C**

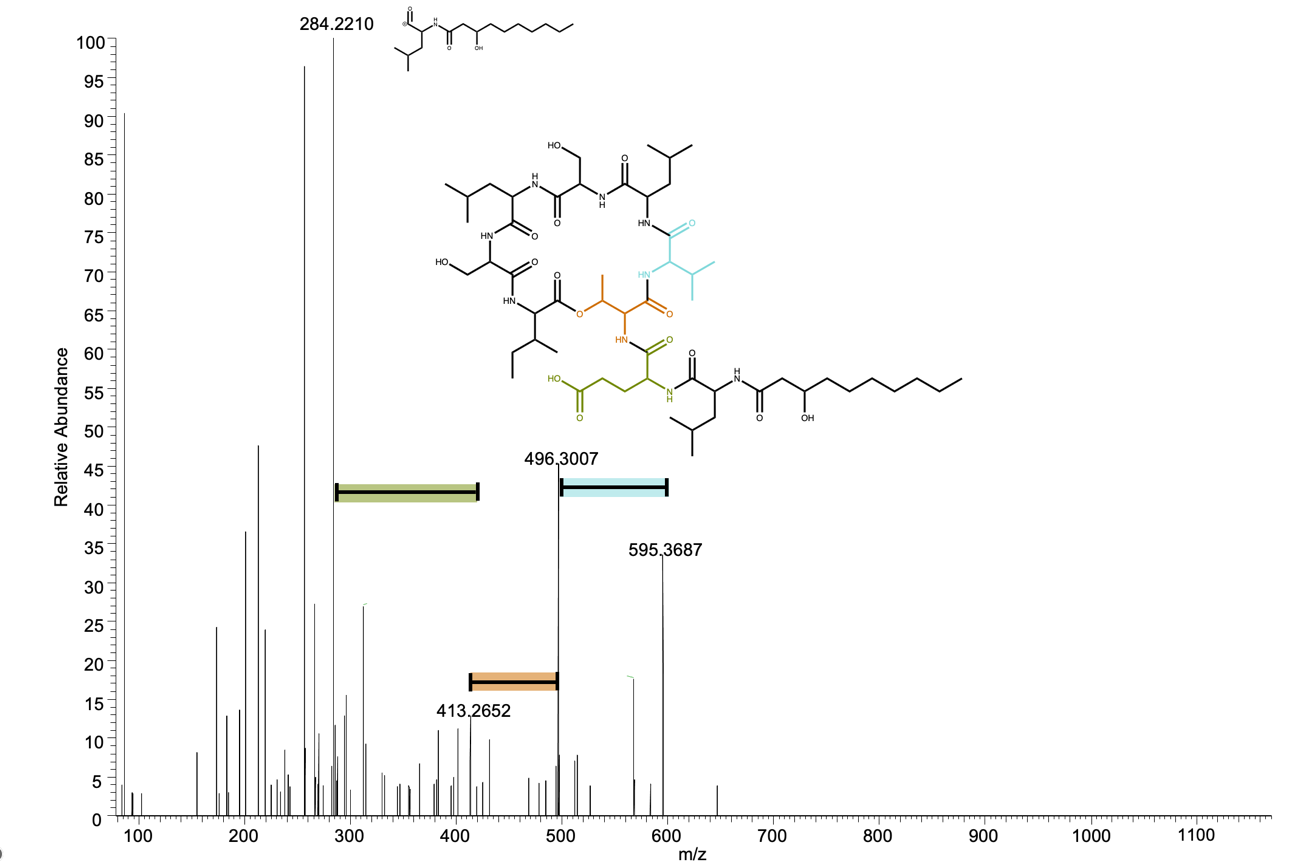

**Figure 7.** HR-LC-ESIMS data of the fraction 10 of the *Pseudomonas* *lurida –* *Legionella* *jordanis* co-culture extract A) Total Ion Chromatogram (TIC) of the fraction 10 of the extracted co-culture. B) Extracted Ion Chromatogram (EIC) (m/z 1126.67 [M+H]) of viscosin. C) Measured fragments and ESI-MS/MS-spectrum of viscosin.
